# Electroactive biofilms on surface functionalized anodes: the anode respiring behavior of a novel electroactive bacterium, *Desulfuromonas acetexigens*

**DOI:** 10.1101/2020.03.04.974261

**Authors:** Krishna P. Katuri, Sirisha Kamireddy, Paul Kavanagh, Ali Mohammad, Peter Ó Conghaile, Amit Kumar, Pascal E. Saikaly, Dónal Leech

## Abstract

Surface chemistry is known to influence the formation, composition and electroactivity of electron-conducting biofilms with however limited information on the variation of microbial composition and electrochemical response during biofilm development to date. Here we present voltammetric, microscopic and microbial community analysis of biofilms formed under fixed applied potential for modified graphite electrodes during early (90 h) and mature (340 h) growth phases. Electrodes modified to introduce hydrophilic groups (−NH_2_, −COOH and −OH) enhance early-stage biofilm formation compared to unmodified or electrodes modified with hydrophobic groups (−C_2_H_5_). In addition, early-stage films formed on hydrophilic electrodes were dominated by the gram-negative sulfur-reducing bacterium *Desulfuromonas acetexigens* while *Geobacter* sp. dominated on −C_2_H_5_ and unmodified electrodes. As biofilms mature, current generation becomes similar, and *D. acetexigens* dominates in all biofilms irrespective of surface chemistry. Electrochemistry of pure culture *D. acetexigens* biofilms reveal that this microbe is capable of forming electroactive biofilms producing considerable current density of > 9 A/m^2^ in a short period of potential induced growth (~19 h followed by inoculation) using acetate as an electron donor. The inability of *D. acetexigens* biofilms to use H_2_ as a sole source electron donor for current generation shows promise for maximizing H_2_ recovery in single-chambered microbial electrolysis cell systems treating wastewaters.

**Highlights:**

- Anode surface chemistry affects the early stage biofilm formation.
- Hydrophilic anode surfaces promote rapid start-up of current generation.
- Certain functionalized anode surfaces enriched the *Desulfuromonas acetexigens*.
- *D. acetexigens* is a novel electroactive bacteria.
- *D. acetexigens* biofilms can produce high current density in a short period of potential induced growth
- *D. acetexigens* has the ability to maximize the H_2_ recovery in MEC.

TOC – Graphical abstract

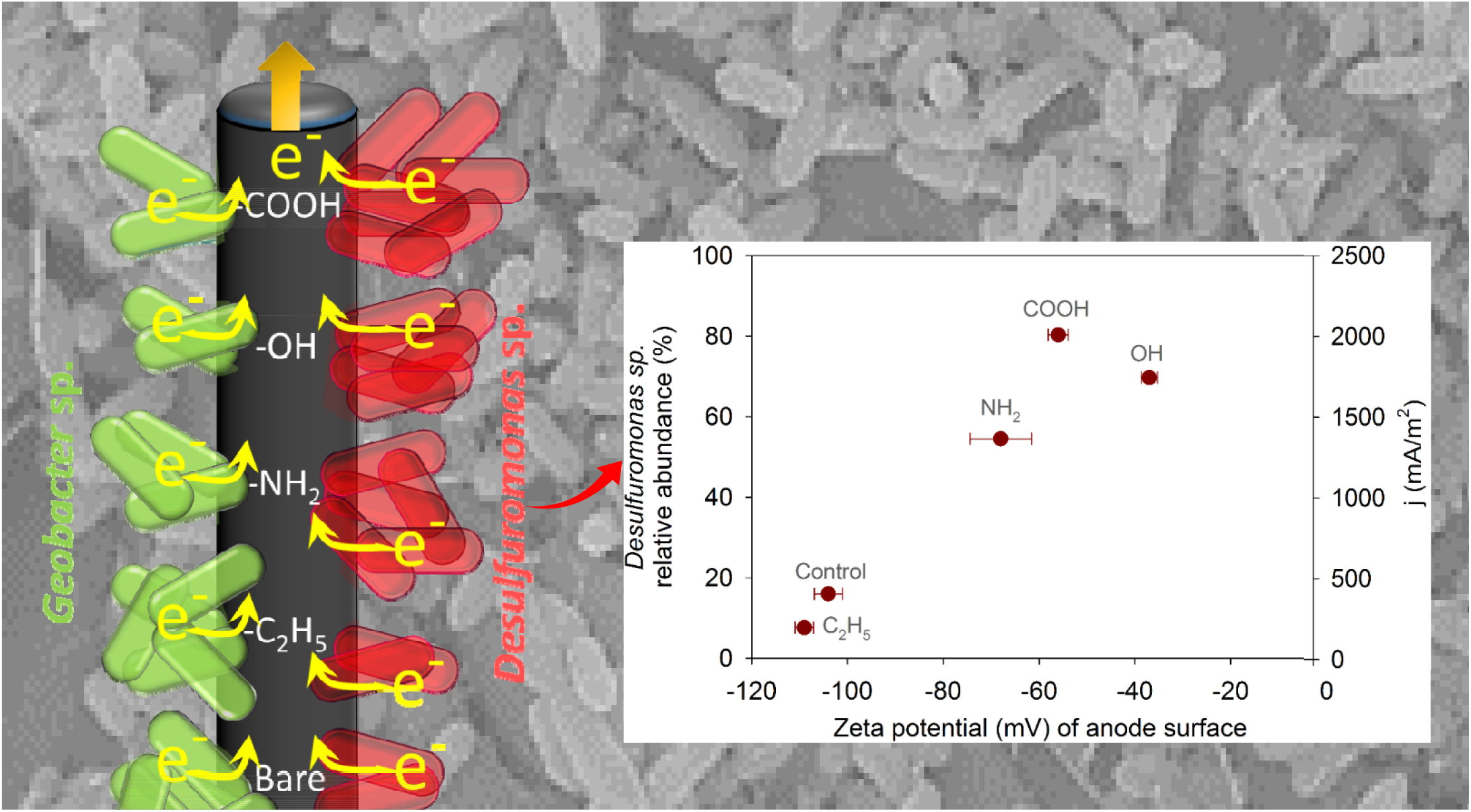

## 1. Introduction

Microbial electrochemical technologies (METs) are electrochemical devices which utilize microbial biofilms formed at a polarized electrode (anode and/or cathode) to drive electrochemical reaction(s) (Rittmann, 2018). An electrochemical potential established at the anode can induce the formation of thick, electron-conducting biofilms composed of special microbial communities known as electroactive bacteria (Schröder et al., 2015). Such biofilms, predominately composed of anaerobic microbes, respire by utilizing an electrode as a terminal electron acceptor in place of natural oxidants such as iron oxide. Potential electroactive bacteria can be found in diverse environments, ranging from the stratosphere (Zhang et al., 2012) to deep Red Sea brine pools/marine sediments (Shehab et al., 2017), including sewage (Patil et al., 2010), sludge, composts, soil, manure, sediments, rumen and agro-industrial wastes (Koch and Harnisch, 2016). Recent study experimentally proved that different known and/or novel electroactive bacteria thrive geographically in a wide range of ecosystems (marshes, lake sediments, saline microbial mats, anaerobic soils, etc.) (Miceli et al., 2012). Thus identifying operational parameters to explore novel electroactive bacteria from mixed culture inoculums with useful metabolic capacities useful for advancing the MET research for niche specific applications. For example, the application of METs has been demonstrated in recovery of bioenergy (bioelectricity and H_2_) from wastewaters (Katuri et al., 2019; Katuri et al., 2018), anoxic NH_4_ removal (Shaw et al., 2019; Vilajeliu-Pons et al., 2018), water reclamation through integration of METs with membrane filtration processes (Katuri et al., 2018; Katuri et al., 2014; Ma et al., 2015; Malaeb et al., 2013), etc. In order to further develop this technology it is imperative to maximize the interaction and to enable efficient electron transfer between the electroactive communities and the electrodes. Understanding the physiology of anodic electroactive bacteria, tuning electrode properties to affect the composition of electroactive bacteria, and tethering and structuring of electroactive communities from different sources at electrodes continues to be a challenge and is the subject of research for the advancement of MES technology.

Several approaches have been developed to establish and improve electrochemical communication between the electroactive bacteria and the anode including chemical treatment of anodes (Dumitru and Scott, 2016), and modification of anode surfaces with mediators (Dumitru and Scott, 2016; Park et al., 2000) and with chemical/functional groups (Artyushkova et al., 2015; Cornejo et al., 2015; Dumitru and Scott, 2016; Guo et al., 2013; Kumar et al., 2013; Lapinsonnière et al., 2013; Picot et al., 2011; Saito et al., 2011; Santoro et al., 2015; Scott et al., 2007). Studies show that the chemical and physical properties of the groups introduced at electrodes can promote or impede electroactive biofilm formation and activity compared to unmodified electrodes, depending on the surface chemistry employed. In general, electrodes modified with charged, hydrophilic functional groups enhanced biofilm attachment, decreased start-up times and improved microbial fuel cell (MFC) or microbial electrolysis cell (MEC) performance (Guo et al., 2013; Kumar et al., 2013; Picot et al., 2011; Saito et al., 2011), whilst the presence of non-polar, hydrophobic groups proved detrimental to biofilm formation and current generation (Guo et al., 2013; Picot et al., 2011).

Most studies on the effect of electrode modification on biofilms focus on current generation at an electrode and on power production in MFC assemblies. Few studies to date investigate the effect of electrode modification on biofilm microbial composition. It has been established that electrode modification can influence microbial composition within biofilms at anodes (Guo et al., 2013; Picot et al., 2011; Santoro et al., 2015). For example, Picot et al., report that biofilms predominately composed of bacteria from *Geobacter* sp., develop on positively charged electrodes (Picot et al., 2011), whereas low cell attachment and *Geobacter* sp. proportion is observed on negatively charged electrodes, with a mixed community evident on neutral electrodes. Guo et al., found abundance of two *Geobacter* sp. (highly similar to *G. psychrophilus* and *G. sulfurreducens*) in matured biofilms (53 day aged) developed on a range of functionalized anode surfaces (Guo et al., 2013). The *Geobacter* relative abundance was found to be higher on anodes functionalized with −N(CH_3_)_3_^+^, −SO_3_^−^ and −OH terminal groups compared to those functionalized with −CH_3_. In addition, a higher relative abundance of *G. psychrophilus* to *G. sulfurreducens* was found in all biofilms, revealing that surface chemistry supported the dominance of electroactive bacteria other than *G. sulfurreducens* (the electroactive bacterium expected to be dominant in anodic biofilms during acetate-fed conditions). However, it should be noted that the inoculum consisted of effluent from the anodic chamber of an existing acetate-fed microbial electrochemical reactor which may be enriched in *Geobacter* species. Using a non-enriched inoculum, Santoro et al., report development of a more diverse consortia consisting of various classes of *Clostridia* and *Proteobacteria* species on functionalized gold electrodes after 45 days (Santoro et al., 2015).

Although such studies provide important insights into the influence of electrode functionalization on microbial community composition, they have been limited to community analysis of consortia in thick biofilms, at the end of relatively long growth periods. Information related to the effect of surface chemistry on the early-stage of microbial biofilm formation, its electromicrobiology and adaptability, is lacking. Here we examine the microbial community composition at modified electrodes (−NH_2_, −COOH, −OH and −C_2_H_5_ terminal groups) for both early (after 90 h growth) and mature (multilayered biofilm after 340 h growth) stage using a non-enriched inoculum, providing insight into biofilm adaptability and maturation. We show that microbial communities can change significantly over time. In addition we present electrochemical characterization of a pure culture of *Desulfuromonas acetexigens*, a gram negative bacterium which was found to dominate in the mature mixed culture biofilms developed at the electrodes using this inoculum.

## 2. Materials and Methods

### 2.1. Electrode preparation

Custom built graphite rod electrodes (0.3 cm diameter, Goodfellow, UK) were prepared by shrouding rod lengths extending out of glass tubes using heat-shrink plastic tubing (Alphawire, UK) and establishing an electrical connection at the rear with a 0.3 cm diameter copper rod (Farnell electronics, Ireland) and silver epoxy adhesive (Radionics, Ireland). The final exposed geometric surface area of the electrode was 3.8 cm^2^. Prior to use these electrodes were sterilized by placement in boiling water for 15 min, and washed several times with distilled water.

Surface functionalization of electrodes to produce −NH_2_, −COOH, −OH and −C_2_H_5_ terminal groups was achieved by electrochemical reduction of the diazonium cation generated *in situ* from the arylamine using either p-phenylenediamine, 3-(4-aminophenyl)propionic acid, 4-aminobenzyl alcohol or 4-ethylaniline, respectively. Briefly, 8 mM of NaNO_2_ was added into a 10 mM acidic solution (0.5 M HCl) of the appropriate arylamine to generate the diazonium cation, followed by electrochemical reduction of the generated aryldiazonium salt by scanning from 0.4 V to −0.4 V vs Ag/AgCl at 20 mV/s for four cycles as described previously (Boland et al., 2008). The resulting modified electrodes were removed and rinsed with large volumes of distilled water, followed by ultrasonication for 1 min to remove any loosely bound species.

### 2.2. Mixed-culture biofilm formation and analysis

The growth medium for forming mixed-culture biofilms was based on *G. sulfurreducens* medium (http://www.dsmz.de, medium no. 826) lacking sodium fumarate and containing 10 mM acetate as electron donor. The medium was purged with N_2_:CO_2_ (80:20) gas mix for 60 min at 10 mL/min gas-flow rate to prepare an oxygen-free solution and then subjected to autoclaving (121 °C, 15 min). After autoclaving, bottles were transferred into an anaerobic glove box (Coy Laboratory, USA) to maintain an anaerobic environment for the medium.

Mixed-culture biofilms were formed by placing electrodes in a custom-built glass electrochemical reactor and application of constant potential (−0.1 V vs Ag/AgCl) using a multi-channel potentiostat (CH Instruments, USA), a common platinum gauze (5 cm × 6 cm) counter electrode and Ag/AgCl reference electrode (3.5 M KCl, BioAnalytical Systems, USA), in the presence of growth medium (500 ml) containing 10 mM acetate as electron donor and 10% of re-suspended granular anaerobic sludge sampled from an internal circulation digester (Carbery Milk Products Ltd., Cork, Ireland) as a mixed-culture inoculum. Prior to inoculation the sludge was crushed and graded by sieving (Ø <0.4 mm) and subsequently concentrated (centrifuge 7000 g, 10 min at 20 °C), washed and re-suspended in 100 ml of sterile de-gassed growth medium. Fresh acetate electron donor, to provide 10 mM concentration, was added to the reactor after 45 h operation. After 90 h, at the end of the batch-feed operation, the reactor was completely drained and electrode samples taken for analysis. The reactor was then filled with fresh growth medium containing 10 mM acetate, with no additional inoculum, and the reactor conditions switched to continuous-mode by pumping culture medium containing 10 mM acetate. Culture medium was maintained in sterile and anaerobic conditions in a reservoir with a working volume of 1 L. The reservoir was equipped with several ports for continuous purging with N_2_, for pumping culture medium into the reactor and for sampling culture medium. Sterile 0.2 mm gas filters were placed on all gas and liquid handling ports except that for pumping the medium from reservoir to reactor. All inoculations were carried out in a sterile anaerobic glove box (Coy Laboratory, USA), and all incubations were performed at 30 °C in a controlled temperature room.

Electrodes sampled after the batch-feed period (90 h growth) were transferred to separate 15 ml vials containing 3 ml of sterile extraction solution (phosphate buffer, pH 7.0, 50 mM). Following biofilm extraction through vigorous vortex, 2 ml of the solution was transferred separately to individual vials for molecular microbial ecology and cell counts analysis. The remainder of the solution (i.e., 1 ml) was filtered through 0.2 µm sterile filter to obtain a cell-free solution for screening of the presence of soluble mediators in the biofilm matrix using CV analysis. A miniature custom-built three-electrode electrochemical cell used to conduct voltammetry in the small-volume electrolyte using a graphite disc (6 mm diameter) and platinum wire as working and counter electrodes, respectively.

In addition, 0.5 cm length of each electrode was sampled and fixed in 2% glutaraldehyde solution for subsequent microscopy analysis. The remaining length of each electrode was transferred to a new electrochemical cell containing fresh growth medium, but with no acetate as electron donor, in order to perform non-turnover voltammetry. A similar sample analysis protocol was adopted for electrodes collected at the end of the continuous-feed growth period (at 340 h).

### 2.3. *D. acetexigens* biofilm formation and analysis

The *D. acetexigens* strain DSM 1397 was cultured at 30 °C in 50 mL air tight, rubber septa-sealed, anaerobic syringe bottles containing 45 mL of growth medium (DSM 148) and subsequently sub-cultured three times (each batch incubated for 3 days) in fumarate-containing growth medium prior to inoculation in the electrochemical cell. The cell pellet collected through centrifugation (at 8000x for 5 min) was used as an inoculum (10% w/v; cell density 3.2 × 10^8^ cells/ml) for the tests.

*D. acetexigens* biofilms were developed on graphite rod (~ 4.8 cm^2^) electrodes by application of constant potential (−0.1 V vs Ag/AgCl) in a three-electrode electrochemical cell configuration using *D. acetoexigens* growth medium (lacking fumarate, resazurin and Na_2_S) as electrolyte with 10 mM sodium acetate as electron donor. Four reactors were operated in parallel in fed-batch mode under the same operational conditions. All inoculations/batch changes were carried out in a sterile anaerobic glove box (Labconco, USA) and incubations were performed at 30 °C in a controlled-temperature room.

The interaction and growth of *D. acetexigens* cells on functionalized (−NH_2_, −COOH, −OH and −C_2_H_5_) anodes during early-stage of growth was studied by placing electrodes in a custom-built glass electrochemical reactor and application of constant potential (−0.1 V vs Ag/AgCl) using a multi-channel potentiostat (CH Instruments, USA), a common platinum gauze (5 cm × 6 cm) counter electrode and Ag/AgCl reference electrode (3.5 M KCl, BioAnalytical Systems, USA), in the presence of *D. acetexigens* growth medium (lacking fumarate, resazurin and Na_2_S) containing 10 mM acetate as an electron donor and 10% w/v inoculum (2.9 × 10^8^ cells/ml), with growth terminated at 25 h after inoculation to measure biomass density on the electrodes.

## 3. Results

### 3.1. Electrode modification

Graphite electrodes were modified through *in situ* formation and subsequent electroreduction of aryldiazonium salts from arylamines as previously described (Boland et al., 2008). Selection of arylamines containing terminal −NH_2_, −COOH, −OH and −C_2_H_5_ functional groups results in formation of surfaces presenting such groups. Voltammograms for the aryldiazonium salt electroreduction process are presented in Fig. S1 showing reduction currents for the salts at −0.13 V for −NH_2_, +0.17 V for −COOH, −0.08 V for −OH, and +0.02 V for −C_2_H_5_ electrodes (V vs Ag/AgCl). The decrease in reduction peak current on the second voltammetric scan is indicative of coupled layer formation (Boland et al., 2008). Zeta potential and contact angle for these electrodes, measured in growth media, are presented in Table S1. Electrodes functionalized to introduce −NH_2_, −COOH and −OH groups display surface zeta potential values that are similar, with the unmodified and −C_2_H_5_ functionalized electrodes showing more negative zeta potentials, in the growth medium. Similarly the unmodified and −C_2_H_5_ functionalized electrodes have the highest contact angles, indicative of surfaces of more hydrophobic character compared to the surfaces functionalized to introduce groups capable of hydrogen-bonding, such as the −NH_2_, −COOH and −OH groups. The cell mats prepared using biofilms of *G. sulfurreducens* or *D. acetexigens* display a relatively low negative zeta potential and low contact angles, indicating material that is hydrophilic and relatively easily wetted.

### 3.2. Electrochemical characterization of biofilms

Induction of growth of electroactive bacteria, and bacterial biofilms, on the electrode surfaces was implemented by polarization of all electrodes (in duplicate) at −0.1 V vs. Ag/AgCl in a three-electrode electrochemical cell configuration using an anaerobic sludge mixed culture inoculum and acetate as carbon and energy source. Initial start-up of bacterial biofilm growth was undertaken in a batch reactor configuration, with removal of inoculum and replenishment of acetate feed at 45 h, when the current started to fall following a period of growth (Fig. 1A). In contrast to other studies (Lapinsonnière et al., 2013) electrode modification did not appear to dramatically affect the time taken for current generation to occur. Onset of a rapid growth in current, for all electrodes, commences between 60-65 h after initial inoculation, which is 20-25 h after removal of inoculum and introduction of fresh acetate as feed in the batch-mode configuration. However, consistent with earlier studies (Cornejo et al., 2015; Guo et al., 2013; Kumar et al., 2013; Santoro et al., 2015), the magnitude of the maximum current density during this early batch-feed cycle is influenced by surface chemistry. Anode surfaces functionalized with chemical functional groups capable of hydrogen-bonding, and therefore more hydrophilic (−NH_2_, −COOH and −OH), yield higher currents compared to the unmodified and −C_2_H_5_ functionalized surfaces (Fig. 1B). Following this initial period (90 h) of electrochemically-induced growth in a batch-fed system a set of electrodes was removed for cyclic voltammetric (CV), microscopic and sequencing analysis. The other set of electrodes was placed in the reactor and continuous flow of acetate-containing cell culture medium commenced at a flow rate of 1 L/day followed by switching to 0.5 L/day after approximately 40 h of continuous feed operation (Fig. 1C, black arrow). During 1 L/day continuous-feed operation all electrodes produced similar current profiles, with the exception of the −C_2_H_5_ functionalized electrode, which produced significantly lower current. Decreasing the flow rate of acetate feed to 0.5 L/day (at ~135 h after initial inoculation of the electrodes, see Fig. 1C) resulted in lower magnitude of current output for all electrodes, except for the −C_2_H_5_ functionalized electrode, which produced now a similar current to that produced by all other electrodes. The decrease in current as a function of flow rate in this study is indicative of acetate mass transport controlled current production. Continuous-feed was maintained up to 340 h after inoculation (Fig. S2), with similar current profiles observed for biofilms developed on all electrodes regardless of flow rate or interruption. Interruption of current generation was implemented to enable recording of CV at several intervals during the continuous-feed period, with these CVs compared to CVs for early-stage biofilms taken at the end of the batch-feed period (90 h).

**Fig. 1.**
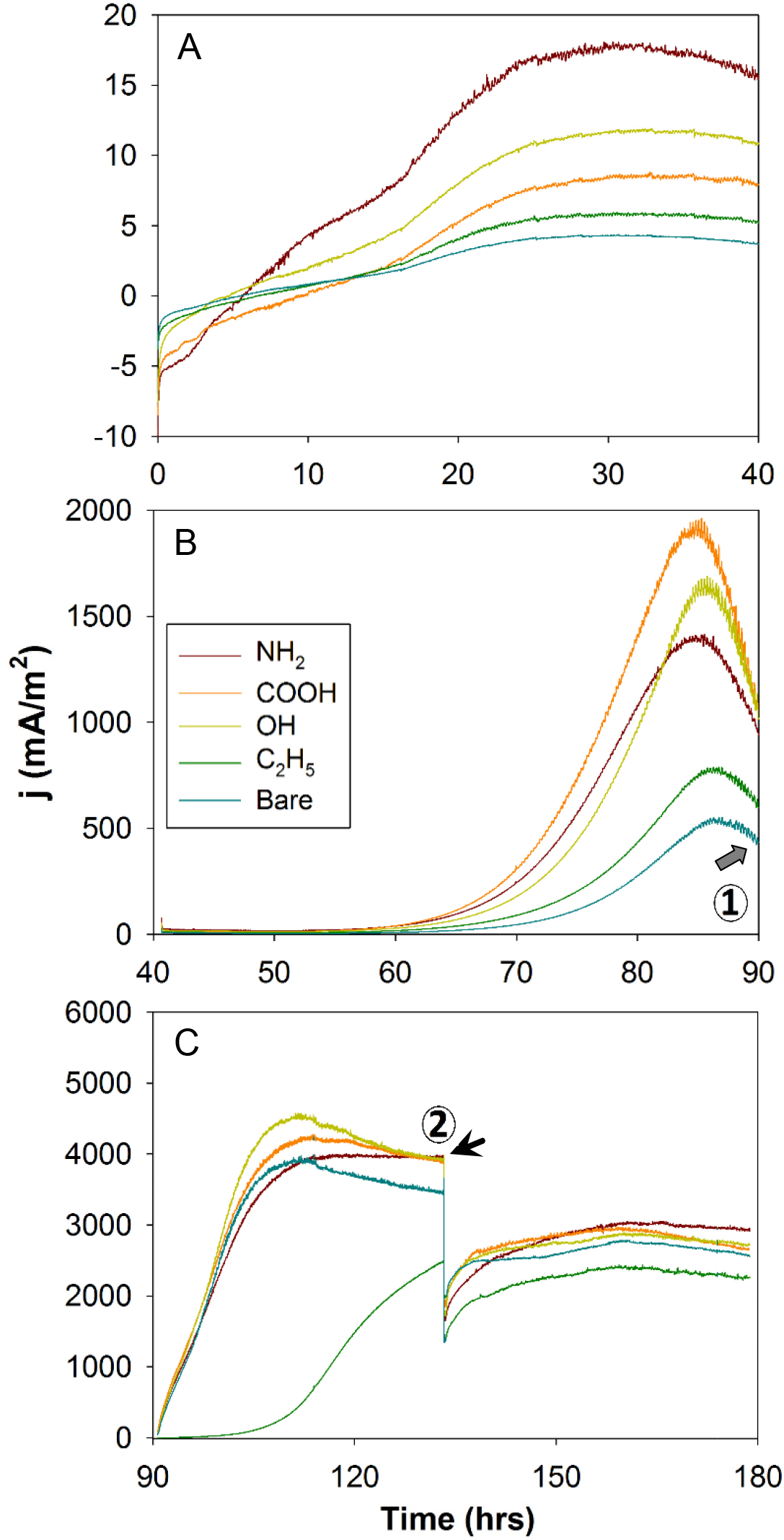
Amperometric response of bare and functionalized graphite electrodes polarized at an applied potential of −0.1 V vs Ag/AgCl. (A), Response for initial start-up, (B), after removal of inoculum and replenishment of acetate and (C), when switching from batch feed to continuous flow feed at a flow rate of 1 L/day, switching to 0.5 L/day at the time indicated by the black arrow. Grey arrow in Figure 1B indicates the time when the biofilms were sampled for SEM and microbial community analysis. Numbers in Figure 1B&C represent the time where CV analysis conducted for the biofilms.

The slow-scan CV response of the early stage biofilms (90 h after initial inoculation) when recorded under substrate-limited conditions all display a well-defined redox couple centered at ~ −0.43 V and an oxidation peak at ~ −0.12 V vs Ag/AgCl, as exemplified by the response obtained at the electrode functionalized to introduce −OH terminal groups (Fig. 2A). The non-turnover analysis of these early-stage biofilms, by transfer into growth medium lacking acetate as electron donor, show three redox responses centered at ~ −0.53 V, −0.36 V and −0.28 V vs Ag/AgCl, shown for the electrode functionalized to introduce −OH terminal groups (Fig. 2A). The CV analysis of the filtered medium harvested from the reactor after 90 h shows a redox couple, with an oxidation peak at ~ −0.13 V (Fig. S3) that is similar to the oxidation peak (~ −0.12 V) observed for the CVs recorded in the growth medium under substrate-limited conditions and to one of the oxidation peaks observed under non-turnover conditions. The slow scan CVs recorded in the presence of 10 mM acetate as electron donor in the electrochemical cell, when flow was halted, show typical sigmoidal shape expected for electrocatalytic oxidation of acetate (Fricke et al., 2008; Katuri et al., 2010; Marsili et al., 2008), as shown for the electrode functionalized to introduce −OH terminal groups (Fig. 2B). There is an increase in the catalytic oxidation current as a function of time after inoculation, despite evidence of uncompensated resistance effect in the CV responses. The sigmoidal shaped CV obtained at 250 h after inoculation (Fig. 2B), when fit to a simple model for steady-state voltammetry (Jana et al., 2014), indicates that electron transfer is dominated by a redox species with an estimated half-wave potential of −0.45 V vs Ag/AgCl, once the approximately 60 Ω uncompensated resistance is accounted for by correcting at each applied potential to achieve the best fit between model and recorded CV. As noted previously (Jana et al., 2014; Torres et al., 2010), this uncompensated resistance is because of cell configuration (distance between working and reference electrode, conductivity of medium, etc.) and probably not a function of low electronic conductivity within the biofilm (Dhar et al., 2017). The half-wave potential of the catalytic CV response, −0.45 V vs Ag/AgCl, correlates well with the potential for the redox couple observed under substrate-limited conditions and the major redox peak under non-turnover conditions (Fig. 2A), while the current density of ~ 4 A/m^2^ is of the same order of magnitude as that observed for multi-layered films of electroactive bacteria on anodes (Jana et al., 2014; Katuri et al., 2012; Marsili et al., 2008).

**Fig. 2.**
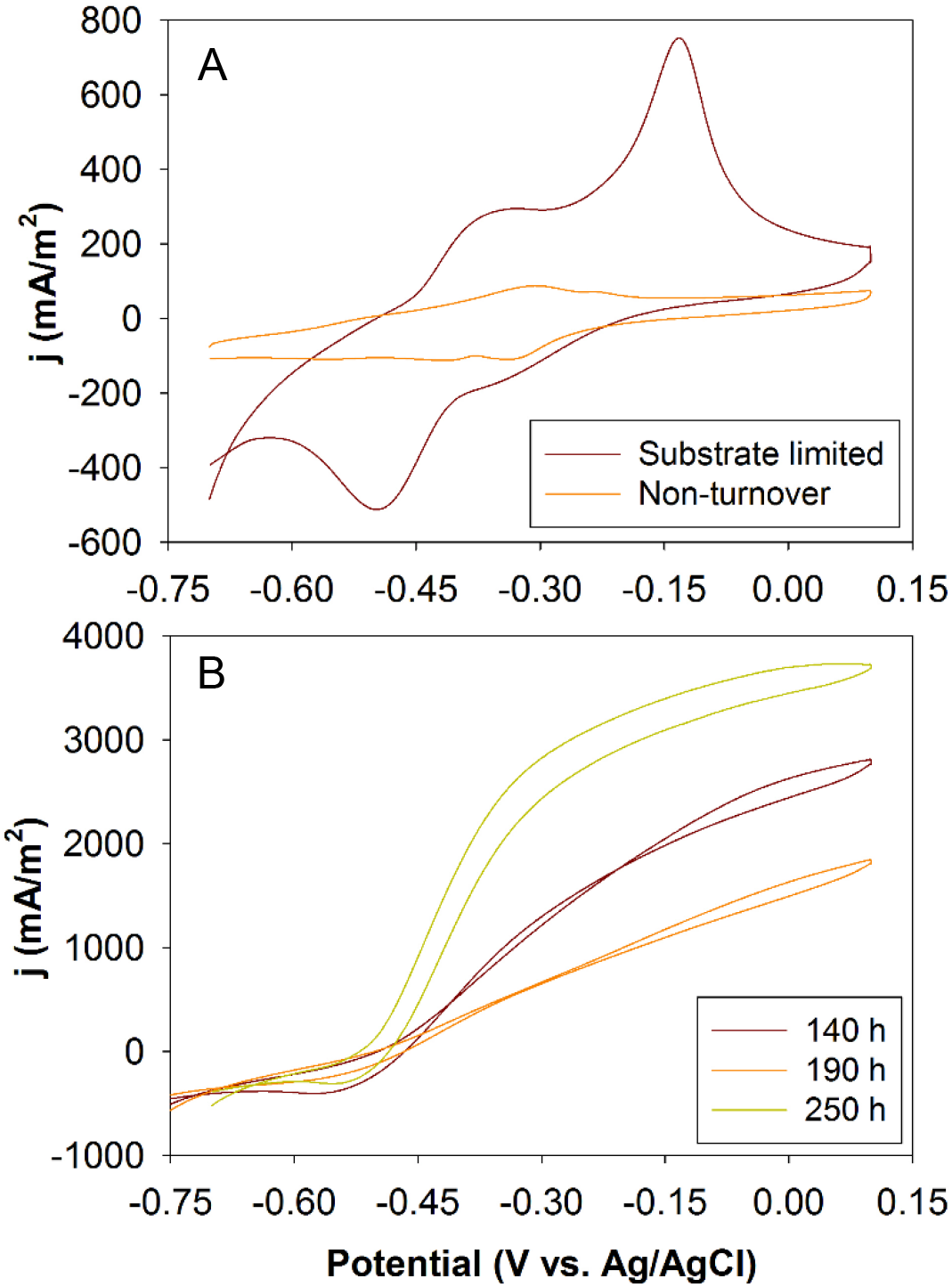
Slow scan CV (1 mV/s) for the −OH functionalized graphite electrodes. (A), Recorded following early-stage growth (90 h) under substrate limiting conditions at the end of the batch feed xand non-turnover conditions in the absence of acetate as electron donor. (B), CV of 140 h, 190 h and 250 h aged biofilms (see Figure S2) in the presence of 10 mM acetate as electron donor.

### 2.3. Microscopy

The SEM images captured at electrodes after early-stage growth (90 h) compared to those captured at a later stage (340 h after initial inoculation) provide additional evidence that the observed amperometric and CV current generation is associated with formation and growth of electrode-attached biofilms. The SEMs after early-stage growth show sparsely and irregularly distributed bacterial cells along with some cell aggregates (Fig. 3A), compared to the presence of thicker and densely-packed biofilms with heterogeneous topography evident in the SEMs of electrodes sampled at 340 h. All the biofilms sampled at 340 h display similar estimated biofilm thickness of ~ 22 µm, with no statistically significant (*P* > 0.05; t test) difference between electrodes, estimated from CLSM imaging (Fig. S4).

**Fig. 3.**
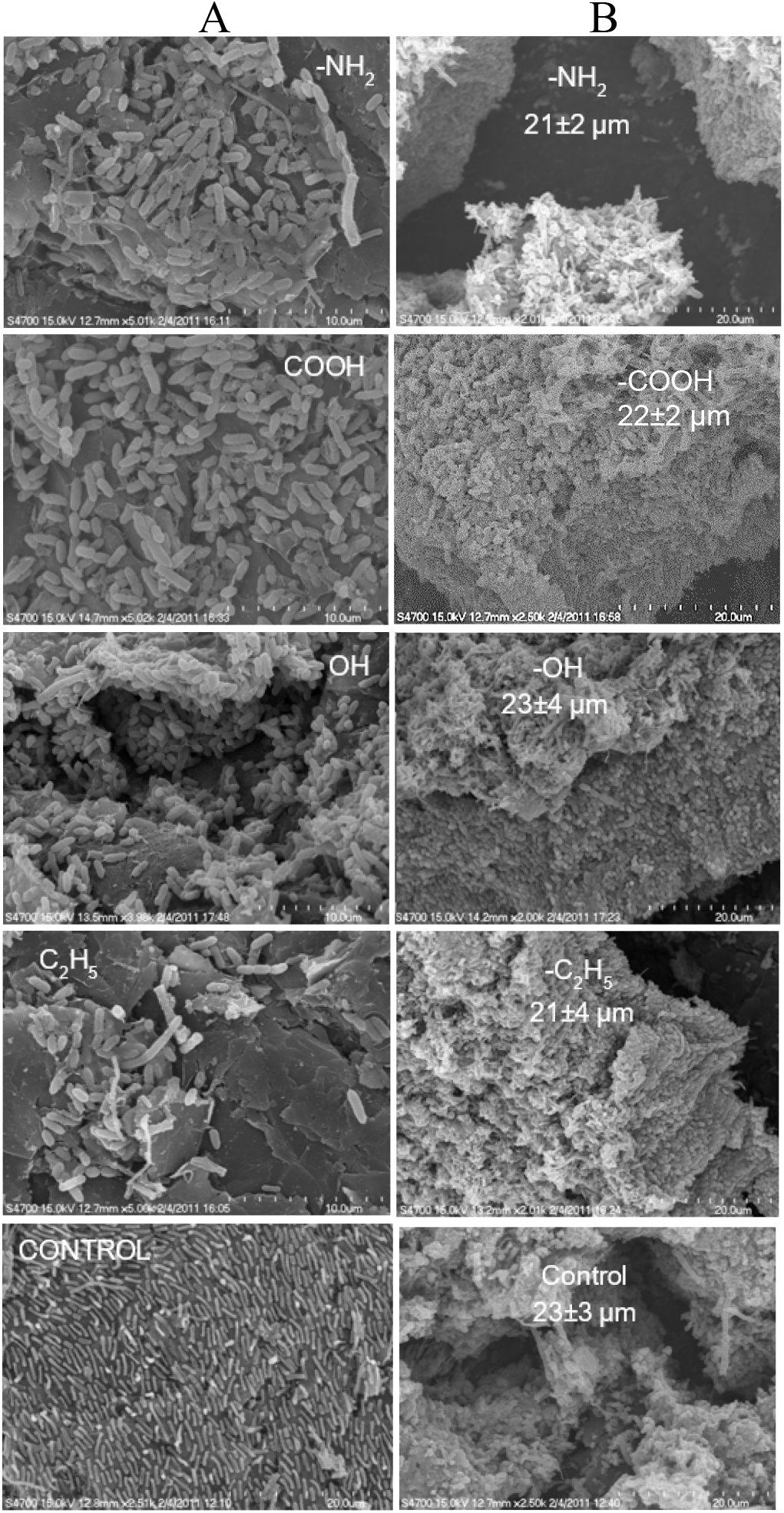
SEM images for biofilm covered control (unmodified) and functionalized graphite electrodes. (A), Biofilms sampled after early-stage (90 h) or (B), later-stage (340 h) biofilm growth conditions (see Fig. 1B and Fig. S2 for details).

### 2.4. Microbial community composition

The early-stage (90 h) and later-stage (340 h) biofilms, as well as the initial anaerobic sludge inoculum, were subjected to 16S rRNA gene sequencing to probe the variation of microbial communities within films prior to, and over, the growth period. Relative abundance of microbes within the biofilms (Fig. 4A) show significant variations as a function of electrode terminal group chemistry and incubation time. The early-stage biofilms have a higher abundance of a genus closely related (99% sequence similarity) to *Desulfuromonas* sp. (dominant OTU), on the electrodes functionalized to introduce −NH_2_, −COOH and −OH groups compared to those of the − C_2_H_5_ functionalized and control (unmodified) electrodes. There is evidence of the presence of known electroactive bacteria i.e., *Geobacter* sp., only for the early stage biofilms grown on −NH_2_, −C_2_H_5_ and control (unmodified) electrodes. Both species were not detected in the inoculum. Selective enrichment of both species and differences in their relative abundance as a function of anode surface chemistry indicates that the anode local environment provides a niche-specific selective pressure for enrichment of a functionally stable bacterial community by growth on the anode surface rather than a random attachment of bacterial cells. For these early-stage biofilms, a clear correlation is evident between current density at the sampling time (90 h, see Fig. 1B) and measured cell density on the anodes (Fig. 4B). In addition there is a clear trend of higher relative abundance of *Desulfuromonas* sp., and current generation, as a function of the estimated zeta potential of the electrodes (Fig. 4C). This observation is supported by the principal components analysis (PCA) of the microbial community in the films (Fig. 4D) showing a clear distinction between the community in the inoculum and the early stage biofilm samples as well as a distinction between hydrophilic surfaces, dominated by *Desulfuromonas* sp., and the unmodified and hydrophobic −C_2_H_5_ surfaces, dominated by *Geobacter* sp. For the later-stage biofilms, sampled after 340 h of reactor operation when the current density is similar for all electrodes, the biofilm composition for all electrodes shifts to become dominated by *Desulfuromonas* sp. (65% – 90%) (Fig. 4A).

**Fig. 4.**
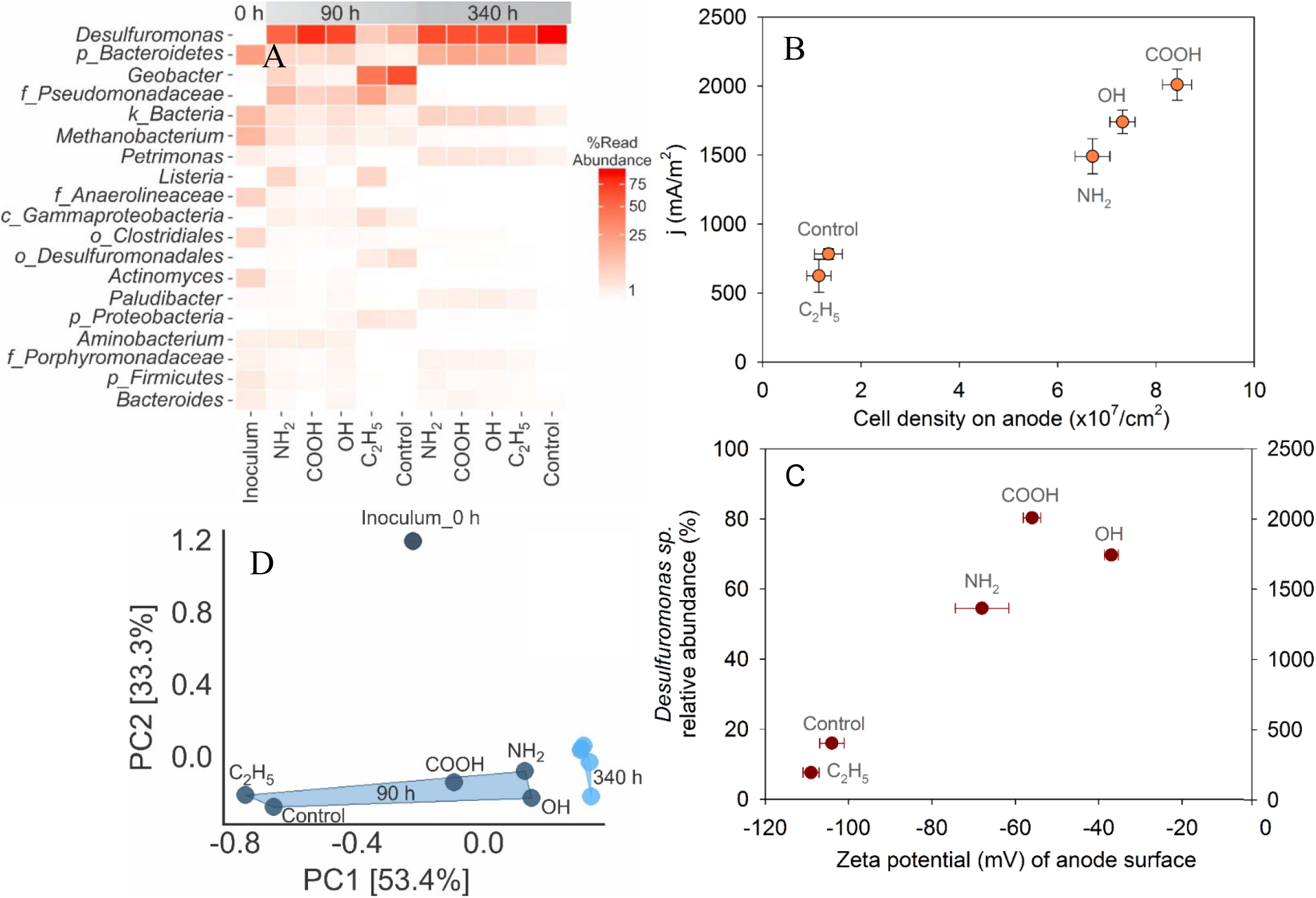
(A), Heat map displaying relative abundance of bacterial reads. The genus level (or lowest taxonomic level possible) relative abundance for inoculum and for biofilms sampled from control (unmodified) and functionalized graphite electrodes after early-stage of batch-feed (90 h) and later-stage of continuous feed (340 h) growth conditions. *k*: kingdom, *p*: phylum, *c*: class, *o*: order and *f* family. (B & C), Influence of anode (unmodified and functionalized graphite electrodes) on biomass growth or zeta potential and its impact on current density after early-stage batch-feed (90 h) growth conditions. Relationship between anode and cell density, and its influence on current production. (B), and correlation between anode surface zeta potential and relative abundance of *Desulfuromonas* sp. and its stimulus on current production (C). (D), PCA analysis showing relationship between biofilm bacterial communities collected over time (90 h and 340 h) and from different electrode (unmodified and functionalized graphite) surfaces. Inoculum sample is also included in the PCA plot.

The remarkable dominance of *Desulfuromonas* sp. prompted further investigation into its role. Subsequent cloning and sequencing of the early-stage biofilm sampled from the electrode functionalized to introduce −COOH terminal groups revealed the dominance of a species closely related (99% sequence similarity) to *Desulfuromonas acetexigens*. Although *D. acetexigens* has been previously identified in electrode-attached biofilms (Ishii et al., 2012; Ketep et al., 2013a) the specific localization of *D. acetexigens* in biofilms and its role in microbial electrochemical systems has yet to be investigated. Our attempts failed to isolate *D. acetexigens* strain from mixed culture biofilms using its natural electron acceptors through both solid/liquid growth approach. Thus, the pure culture of *D. acetexigens* (DSM 1397) purchased from DSMZ was used for conducting the electromicrobiology experiments.

Induction of growth of *D. acetexigens* bacterial biofilms on an unmodified graphite rod electrode surface was implemented by polarization of electrodes at −0.1 V vs. Ag/AgCl in three-electrode electrochemical cell configuration using *D. acetexigens* culture as inoculum and batch-feeding with acetate as substrate and energy source in an appropriate cell culture medium (see experimental details). The evolution of current over time, in this reactor, is similar to that observed for other pure culture electroactive bacteria under a continuous applied potential, such as *G. sulfurreducens* (Fricke et al., 2008; Jana et al., 2014; Katuri et al., 2010; Liu et al., 2008; Marsili et al., 2008) i.e., cycles of a rapid rise in current when acetate is introduced and then a relatively sharp fall as a consequence of acetate substrate depletion, as shown in Fig. 5A. Remarkably, relatively rapid initial current is observed without a substantial lag phase during the first batch of operation with a peak in the current density of 9.2 ± 0.4 A/m^2^ obtained only 19.3 ±0.4 h following initial inoculation into the reactor. Little significant further improvement to the maximum current density during the batch-feed cycles is observed, with peak current density reaching a maximum of ~10 A/m^2^ over the ~210 h growth period in the reactor.

**Fig. 5.**
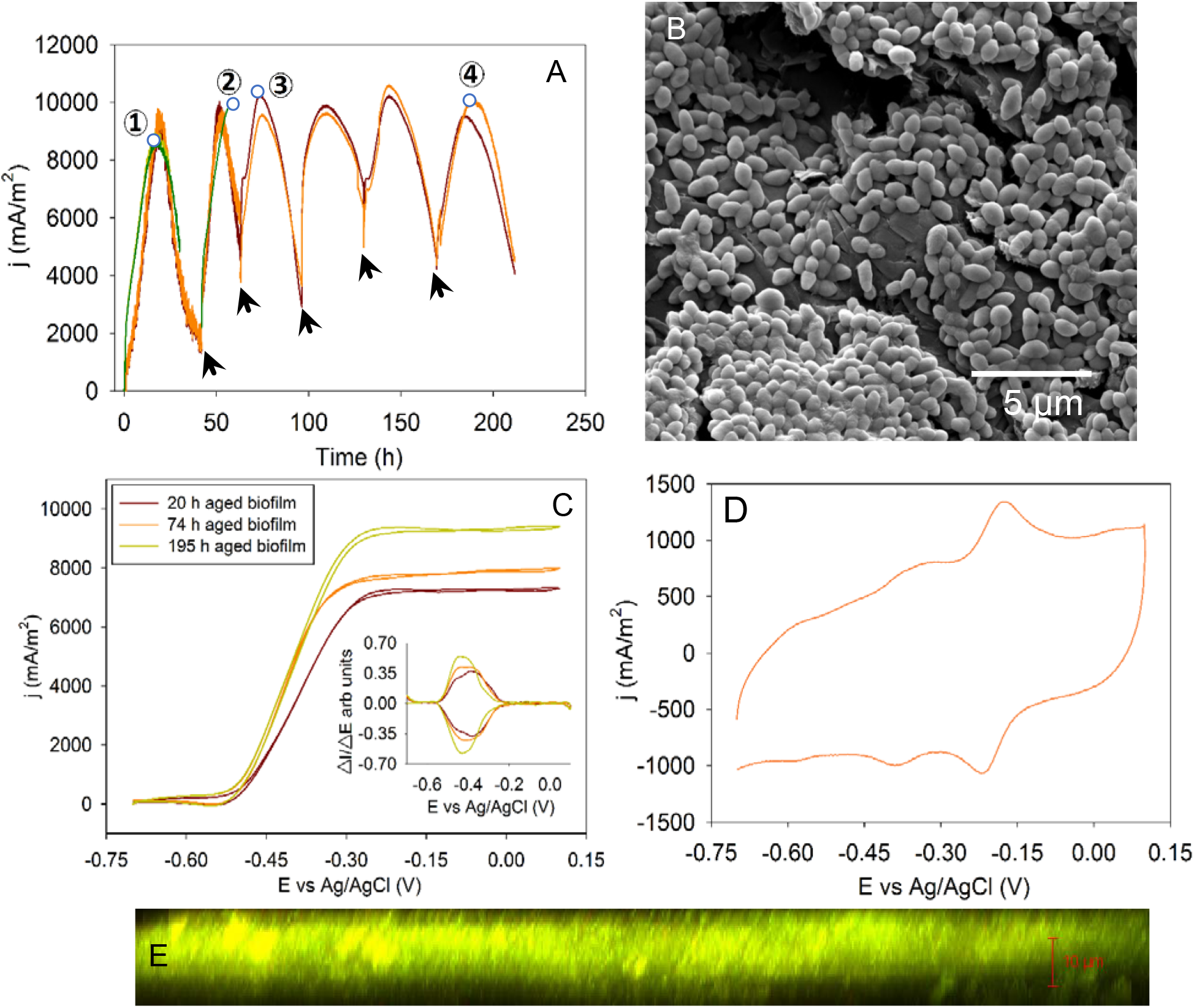
Electrochemical behavior of *D. acetexigens* biofilms. (A), Amperometric response of graphite rod electrodes in the electrochemical reactor, at an applied potential of −0.1 V vs Ag/AgCl, during batch-feed operation, where the arrows represent change of feed. (B), SEM image of electrode sampled at time indicated by (1) in (A). (C), *In-situ* CVs (1 mV/s) conducted at times indicated by (1), (3) and (4) in (A), with inset representing the first derivative of the CVs. (D), Non-turnover CV (1 mV/s) recorded in anaerobic phosphate buffer (100 mM, pH 7.0) for the electrode sampled 210 h after inoculation. (E), CLSM image of ~ 60 h aged biofilm sampled at time indicated by (2) in (A).

*In-situ* recording of slow-scan CVs at specific intervals (20 h, 74 h and 195 h) after initial inoculation provides the characteristic sigmoidal shape, indicative of microbial-electrocatalytic oxidation of acetate substrate by a *D. acetexigens* biofilm on the electrode surface (Fig. 5B), as observed for the mixed-culture biofilms. Examination of the first derivative of the CVs indicates the presence of a dominant redox transition with a half-wave potential of approximately −0.42 V vs Ag/AgCl (inset of Fig. 5C). The CVs in the presence of acetate show an increase in steady-state currents in progressing from the early-stage (20 h) biofilm to those recorded at later stages of growth (74 h and 195 h), an increase that is also observed in the fixed potential amperometric response at those sampling times (Fig. 5A). The non-turnover analysis of the later-stage biofilm (210 h after inoculation), by transfer into pH 7.0 phosphate buffer electrolyte lacking acetate as electron donor, shows three clear redox responses centered at ~ −0.58 V, −0.37 V and −0.20 V vs Ag/AgCl (Fig. 5D). No discernible redox response is observed in CVs recorded for the reactor bulk liquid.

The catalytic activity of *D. acetexigens* with formate or H_2_ (intermediates of anaerobic digestion process) as an electron donor was tested separately in a three-electrode electrochemical cell under −0.1 V vs. Ag/AgCl fixed anode potential. A maximum current density of 4.4 ± 0.3 A/m^2^ was generated over a growth period of 26 h following inoculation (Fig. 6) using formate as electron donor. When the reactor feed was altered to include acetate as an electron donor instead of formate, maximum current density of 10.6 mA/m^2^ over a short period of reactor operation resulted. In a parallel experiment current generation practically ceased when feed was altered to include H_2_ as an electron donor instead of formate. Also no H_2_ consumption is observed during the test period. A similar behavior is observed for biofilms developed initially using acetate which is then altered to H_2_ as the electron donor (data not shown).

**Fig. 6.**
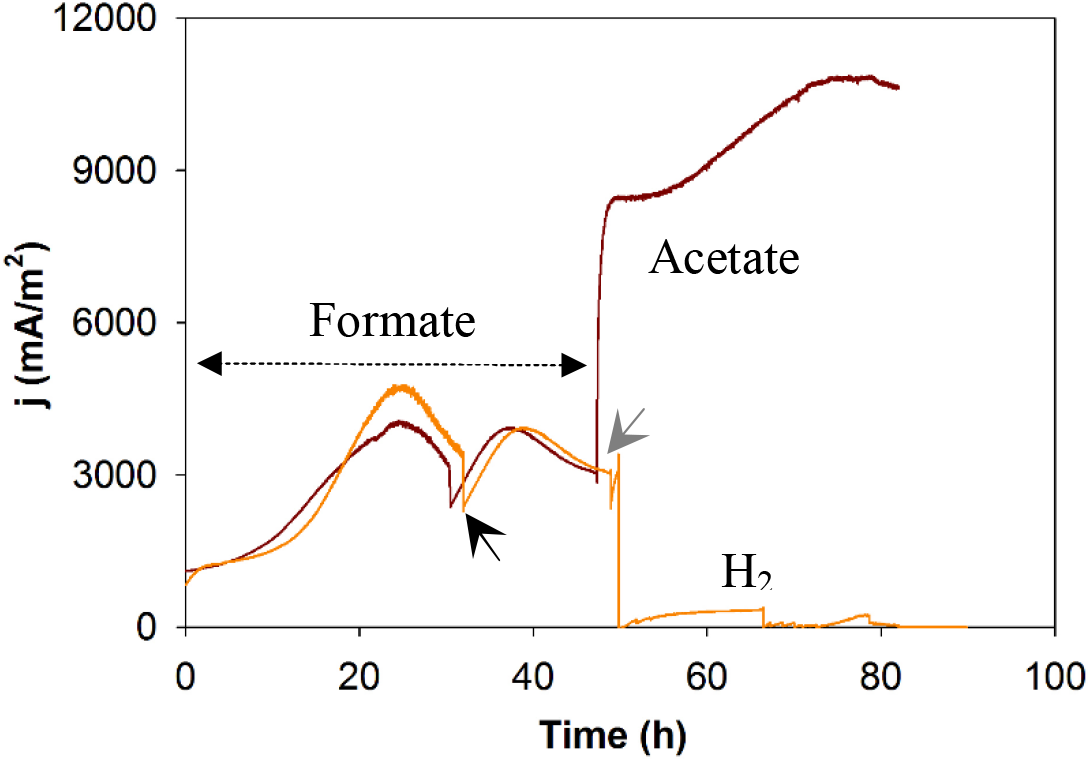
Amperometric response of *D. acetexigens* biofilms grown on graphite rod electrodes at an applied potential of −0.1 V vs Ag/AgCl in the electrochemical reactor during batch-feed operation. Black arrows represent change of feed. Gray represents the time when respective biofilms switched to acetate or H_2_ as an electron donor instead of formate. The same electron equivalent substrate concentration (i.e., 20 mM formate and 5 mM acetate) is used for the tests.

## 4. Discussion

Electroactive bacteria attachment and biofilm formation is considered as a primary step in the microbial-electrode enrichment process. The key selective pressure for this electricigen enrichment in MES is extracellular electron transfer (EET) with the anode acting as an electron acceptor. The EET in electroactive biofilms is proposed to occur through production of exogenous mediators by the biofilms or through self-exchange between outer-membrane bound c-type cytochromes present on certain microbial cell surfaces facilitating electron transport through the film to the solid electrode surface and between cells at the interface between the biofilm and the solid-state anode (Nevin et al., 2009), with some postulating EET occurring by electronic conduction along structured protein channels (pili) (Sure et al., 2016). However, the crucial factors that control initial electrochemical current generation by interaction between the external bacterial cell surface and solid anodes is not yet clearly elucidated. Achieving insight into conditions that control this interaction is therefore crucial for shaping the anodic microbial community and improving MES technology.

Results presented here demonstrate that terminal group chemistry on graphite electrodes influences the microbial community composition and relative abundance during the early-stage of biofilm formation and growth, (Fig. 1B and Fig. 4), for biofilms grown under fixed applied potential in a single chamber electrochemical cell using an inoculum harvested from an anaerobic digester treating dairy plant wastewaters, confirming observations by others using a range of inocula and conditions (Guo et al., 2013; Picot et al., 2011; Santoro et al., 2015). The majority of electroactive bacteria reported to be present in anodic biofilms are gram negative (Read et al., 2010), possessing negatively charged bacterial cell surfaces (Santoro et al., 2015). Thus, positively charged electrode surfaces are thought to promote strong electrostatic interactions between the electrode surface and the negatively charged electroactive bacteria (Guo et al., 2013; Kumar et al., 2013; Lapinsonnière et al., 2013; Picot et al., 2011; Santoro et al., 2015). However, Guo et al., report that start-up of current generation is more rapid on glassy carbon electrodes functionalized to introduce hydrophilic (−N(CH_3_)_3_^+^, −OH and −SO_3_) groups, regardless of the charge on the functional group (Guo et al., 2013), compared to start-up on −CH_3_ terminated surfaces, and this was confirmed by Santoro et al., using self-assembled monolayers on gold (Santini et al., 2015). We find that graphite electrodes functionalized to introduce −NH_2_, −COOH and −OH display higher currents during initial stage of biofilm growth, under batch-feeding of acetate as electron donor, compared to electrodes functionalized to introduce −C_2_H_5_ and unmodified graphite electrodes, under the same operating conditions (Fig. 1 & Fig. 4 B&C). The capacity to permit cell growth, and to generate current is clearly related to the surface charge on the electrodes, as represented by the zeta potential measured in the growth medium (Fig. 4C), with the −C_2_H_5_ and bare electrodes displaying the more negative zeta potentials. It does not appear that the current generation is related to the sign of the charge on the surface terminal group, as the-NH_2_ and −OH groups are expected to be neutral while the −COOH groups are expected to be de-protonated and negative under the cell culture medium conditions (pH 6.8). The ability to promote preferential electroactive bacteria attachment during the initial phase of colonization may therefore be through capacity to interact electrostatically, for example through formation of hydrogen bonds, with the bacterial cell surface, noting that the dipole moment of each of aniline, phenylpropionic acid and benzylalcohol, presumed to be the dominant terminal molecules at the −NH_2_ and −COOH and −OH functionalized electrodes, is above 1.5 D while the dipole moment for ethylbenzene, present at the −C_2_H_5_ functionalized electrode, is 0.58 D (Ray, 2017). It has been highlighted that *Shewanella loihica PV-4* has capability to generate five-fold higher current on a hydrophilic compared to that on a hydrophobic electrode under fixed anode potential growth conditions (Ding et al., 2015). Thus local polarity is a crucial factor in inducing preferential colonization of electrodes by electroactive bacteria, enhancing current during the early-stage of electroactive biofilm growth.

The voltammetric analysis at low, or absent, acetate levels, for early stage biofilms show a redox signal centered at potential of −0.43 V vs Ag/AgCl similar to that observed for biofilms induced to grow from pure culture of *G. sulfurreducens* (Fricke et al., 2008; Katuri et al., 2010; Marsili et al., 2008) or *G. anodireducens* (Sun et al., 2014), as well as a signal at ~ −0.2 V vs Ag/AgCl that resembles the redox peak observed with the filtered biofilm-extracted solution, indicating that the biofilms produce an extracellular, water soluble, mediator. The redox potential of the detected mediator is comparable to that of phenazine-type redox mediators secreted by a range of microbes (Wang et al., 2010), including *Pseudomonadaceae* microbes, which have been detected to be present in the biofilms (Fig. 4A).

The effect of experimental conditions such as inoculum selection, anode surface charge (Guo et al., 2013), electrode nano/micro-scale topography (Champigneux et al., 2018) and cell surface polarizability (Wang et al., 2019) on the microbial composition in electroactive biofilms is not as yet widely understood. For example, Guo et al., studied the microbial community composition of anodic biofilms developed on a range of surface functionalized glassy carbon electrodes (−N(CH_3_)_3_^+^, −OH, −SO_3_ and −CH_3_) and found predominance of *Geobacter* sp., in the biofilms after 52 days of operation, irrespective of functionalized anode tested, but with a lower predominance in −CH_3_ functionalized electrodes (Guo et al., 2013). This is presumably because of seeding of reactor with effluent from an actively operating (more than a year) acetate-fed microbial electrochemical system. However, in our study, noticeable differences in the relative abundance of *Desulfuromonas* sp., (−COOH > −OH > −NH_2_ > Control > −C_2_H_5_) and *Geobacter* sp., (Control >−C_2_H_5_ > −NH_2_ > −COOH > −OH) is found for the early stage anodic biofilms formed on functionalized electrodes (Fig. 4A) suggesting that the micro-environment (i.e., polarity, hydrophilicity, charge, etc.) can stimulate adhesion as well as viability of *Desulfuromonas* sp., over that of *Geobacter* sp.

When the biofilms are matured, over the 340 h growth period, there is greater similarity in the communities detected within the films (Fig. 4A & D), as well as in the currents generated (Fig. 1) due to convergence of biofilm microbial composition predominantly to *Desulfuromonas* sp., This result strongly signifies that a single bacterial genus, i.e., *Desulfuromonas* sp., is the major contributor to current generation in these biofilms. Subsequent cloning and sequencing of the early-stage biofilm sampled from the electrode functionalized to introduce −COOH groups revealed the dominant species as *D. acetexigens*. Colonization by *D. acetexigens* in the early stage of biofilm formation could limit species diversity within the biofilms over the growth period by competing for the space and electron donor on the polarized anode surfaces perhaps because of a superior anode-respiring capability over *Geobacter* sp. A similar trend was reported by Ishii et al., with a shift in dominance from *Geobacter* sp. towards a phylotype closely related to *D. acetexigens* in the anodic biofilm after long term operation (> 200 days) of single-chamber, air-cathode MFCs using primary clarifier effluent as a source of feed and inoculum, and carbon cloth as anode (Ishii et al., 2012). The same team (Ishii et al., 2014) had reported that the relative abundance of *D. acetexigens* over *Geobacter* sp. in an anodic biofilm increased over 3 months of acetate feed operation in a MFC inoculated with a sediment slurry from a lagoon.

*D. acetexigens* is a gram-negative bacterium belonging to the family *Desulfuromonadaceae* (class *Deltaproteobacteria*). It has been detected in freshwater sediments and digester sludge of wastewater treatment plants, and links acetate oxidation to sulfur reduction (Finster et al., 1994). The existence of *Desulfuromonas* sp., at low relative abundance has been reported in the anodic biofilms where domestic sewage (Ishii et al., 2012; Ketep et al., 2013a), raw paper mill effluents (Ketep et al., 2013a; b) and lagoon sediment (Ishii et al., 2014) were used as the source of inoculum. There have been no reports, to date, characterizing the performance of biofilms of pure *D. acetexigens* induced to grow on electrodes. The electrochemical activity of *D. acetexigens* films induced to grow from a pure culture inoculum on graphite electrodes is reported here (Fig. 5A&C). The amperometric and voltammetric signals confirm that *D. acetexigens* has the ability to use a graphite anode as an electron acceptor in the presence of acetate as electron donor, respiring on the anode, providing evidence that this bacterium is responsible for such signals observed for biofilms in previous reports (Ishii et al., 2012; Ishii et al., 2014; Ketep et al., 2013a; b). The slow-scan voltammetry of biofilms in the presence of acetate display a half-wave redox potential of −0.42 V vs Ag/AgCl comparable to that observed for the biofilms grown using the anaerobic sludge as inoculum on the graphite electrode (Fig. 2) and to redox potentials reported for membrane bound redox proteins expressed by other electroactive bacteria (Katuri et al., 2010). Non-turnover voltammetry of *D. acetexigens* biofilms (Fig. 5D) show redox peaks centered at ~ −0.58 V, −0.37 V and −0.20 V vs Ag/AgCl representing presence of a number of redox moieties at potentials comparable to those reported for biofilms of *G. sulfurreducens* (Fricke et al., 2008; Katuri et al., 2012). Remarkably, rapid onset of current to generate ~ 9 A/m^2^ in less than 20 h after start-up is observed with formation of a thin layer of adhered cells (Fig. 5B). However, further improvement in biofilm growth followed by fed-batch operation did not significantly improve the magnitude of the current generation. A current density of ~ 9.7 A/m^2^ is observed for a ~ 60 h aged biofilm having a thickness of ~10 µm (Fig. 5E). Although comparisons are difficult because of different conditions (cell configuration, electrodes, medium, inoculum and its cell density and growth phase etc.), the start-up of *D. acetexigens* for current generation is faster than using pure cultures of *G. sulfurreducens*. For example, for *G. sulfurreducens* current density of up to ~ 5 A/m^2^ is achieved after 72 hours of growth at an applied potential of 0 V vs. Ag/AgCl at graphite electrodes using a high proportion of inoculum in a batch feed mode (Marsili et al., 2008), while a current density of ~ 9 A/m^2^ is generated only after 142 h (Katuri et al., 2012) or 85 h (Jana et al., 2014) of repeated batch mode experiments at an applied potential of 0 V vs. Ag/AgCl at graphite electrodes.

The current generation using formate as electron donor (Fig. 6) confirms that *D. acetexigens* can link formate oxidation with anode respiration. However, a 2.5 fold increase in current density generation achieved for *D. acetexigens* biofilms when formate is replaced with the same electron equivalent acetate concentration suggests that *D. acetexigens* biofilms are more active with acetate as the electron donor. Recycling of H_2_ as the electron donor by *G. sulfurreducens*, a well-studied electroactive bacteria, can adversely affect the energy harnessed in a single-chambered MEC. The inability of *D. acetexigens* biofilms to use H_2_ for current generation provides opportunity to maximize recovery of wastewater energy as H_2_ using MET.

## 5. Conclusion

Results show a clear influence of functionalization of anodes on primary adhesion of microbes during the early stage of biofilm formation for electricigenesis, providing differences in onset of current and time to achieve maximum, steady-state, current. Although we do not yet know whether functionalized electrodes can be used to differentially stimulate enrichment of one electroactive bacterium over another, or rate of anodic biofilm formation from different inoculum sources, the results obtained in this study using anaerobic sludge as inoculum suggest that biomass adhesion as well as its activity, biofilm morphology (i.e. spatial distribution of cells) and relative abundance of electroactive bacteria during the early stage of biofilm formation are clearly affected by the anode surface characteristics. More importantly, the hydrophilic surfaces promote rapid start-up of current generation. Although the underlying mechanism is unclear, anodes functionalized with hydrophilic terminal groups result in the enrichment of a *Desulfuromonas* sp. in the biofilm compared to initial dominance of *Geobacter* sp. on more hydrophobic electrodes. This *Desulfuromonas* sp. is identified to be *D. acetexigens*, and the study of early-stage biofilms of this species is presented confirming its electroactive response. Superior electrocatalytic performance with distinct anode respiring properties of *D. acetexigens* biofilms will prove advantageous for microbial-anode performance in METs. Identification of selection pressure to increase the abundance of *D. acetexigens* in anodic biofilms (perhaps through bio-augmentation or identifying the optimal differential growth conditions) can maximize resource recovery from the wastes. For example, metabolic activities of *D. acetexigens* biofilms (such as efficient EET properties and absence of H_2_ recycling as an electron donor) will favor maximum recovery of energy as H_2_ in single-chambered MEC systems for wastewater treatment.

## Acknowledgments

This work was supported by the Center Competitive Funding Program (Grant No. FCC/1/1971-05-01) and the Competitive Research Grant (URF/1/2985-01-01) from King Abdullah University of Science and Technology (KAUST) and by a Charles Parsons Energy Research Award, through Science Foundation Ireland.

