## Supplemental for "Electroactive biofilms on surface functionalized anodes: the anode respiring behavior of a novel electroactive bacterium, *Desulfuromonas acetexigens*"

**DNA extraction, PCR, and 16S rRNA gene sequencing**

Mixed-culture biofilms sampled from electrodes after batch-feed (90 h) and continuous-feed (340 h) growth were subjected to genomic DNA isolation using the PowerSoil DNA extraction kit (MO BIO Laboratories, Inc., Carlsbad, CA) following the manufacturer's instructions. Triplicate PCR reactions were performed for each sample in a 25 µL reaction volume using the HotStarTaqPlus Master Mix (Qiagen, Valencia, CA), 0.5 µM of each primer, and 100-200 ng of template DNA. Primers targeting the Archaea and Bacteria, 16S rRNA gene region V4: [515F] GTGYCAGCMGCCGCGGTA and [805R] GACTACHVGGGTATCTAATCC used for amplifying 16S rRNA genes (Ye et al. 2016). PCR was performed using a C1000 Thermal Cycler (BioRad, Hercules, CA) with initial denaturation at 95 °C for 5 min, followed by 27 cycles of denaturation at 94 °C for 1 min, annealing at 56 °C for 1 min, extension at 72 °C for 1 min and final extension at 72 °C for 7 min. The triplicate PCR products from each sample were pooled and then purified using the Qiaquick gel extraction kit (Qiagen, Valencia, CA) according to the manufacturer's protocol. The concentration of the PCR products was measured with a Qubit® 2.0 Fluorometer using the PicoGreen® dsDNA quantitation assay (Invitrogen, Carlsbad, CA). The purified barcoded amplicons from each sample were pooled in an equimolar concentration and sequenced on the Roche 454 FLX Titanium genome sequencer (Roche, Indianapolis, IN) according to manufacturer's instructions.

The 16S rRNA gene sequences were processed using the Quantitative Insights Into Microbial Ecology (QIIME v 1.7.0) pipeline (Caporaso et al. 2010). Raw reads were first de-multiplexed, trimmed and filtered for quality. The minimum acceptable length was set to 200 bp (Caporaso et al. 2010). Sequences were clustered into operational taxonomic units (OTUs) at 97% sequence similarity using the uclust algorithm (Edgar 2010). A representative sequence from each OTU was aligned using PyNAST (Caporaso et al. 2010), and these were phylogenetically assigned to a taxonomic identity (genus level) using the RDP Naive Bayesian rRNA classifier at a confidence threshold of 80% (Wang et al. 2007). Chimeric sequences were identified and removed from the aligned sequences using chimera Slayer as implemented in QIIME. Ten most abundant OTUs from each sample were further classified at the species level using the BLAST database (http://blast.ncbi.nlm.nih.gov/Blast.cgi). Principal component analysis (PCA) was used to compare the similarity between the different electrode samples using the statistical software PRIMER 6 (version 6.1.13) and PERMANOVA+ add on (version 1.0.3) and a PCA plot was generated based on Bray–Curtis dissimilarity index (Bray and Curtis 1957) .

**PCR cloning and sequencing**

The PCR product of the sample obtained after batch-feed growth (90 h) from the -COOH functionalized electrode was used for cloning. Universal primers (27F and 1492R) used for the 16S rRNA PCR amplification. The purified PCR fragments were ligated into a pEASY-T cloning vector and cloned into Trans1 chemically competent cells according to the manufacturer’s instructions (TransGen, China). Transformants were screened by blue/white colony selection on agar containing X-gal/IPTG and 100 μg/mL ampicillin. White colonies were randomly selected and grown overnight in 3 mL LB medium containing 100 μg/mL ampicillin. Plasmids were isolated using a plasmid purification kit (Biomiga). The insert in the plasmid was checked by PCR using primers M13F and M13R as previously described (Messing 1983). Twenty-two positive clones were selected and submitted for 16S rRNA gene sequencing (454 FLX Titanium, Roche, Indianapolis, IN). Clone library coverage was calculated by Good’s coverage (1-n/N)×100, where n is the number of single reads and N is the number of total reads.

**Surface charge and contact angle measurement**

A zeta potential analyzer (Anton Paar, Austria) and goniometer CAM200 (KSV, Finland) were used to determine the surface charge and contact angle of the graphite electrodes, and cell-mats of *G. sulfurreducens* and *D. acetexigens* on filter paper.

To prepare the cell-mats, *G. sulfurreducens* (DSM 12127, from German Collection of Microorganisms and Cell Culture Centre) or *D. acetexigens* (DSM 1397) were sub-cultured repetitively in 100 mL air tight, rubber septa sealed, anaerobic syringe bottles containing 70 mL of growth medium. The cells (200 ml) were harvested by centrifugation (8000g, 10 min) and washed twice with saline solution (0.9% NaCl). The collected biomass pellets were re-suspended separately in distilled water and vortexed before filtration through micropore filters (0.2 µm, Whatman) which were subsequently dried at 80 °C for 1 h to remove moisture from films in order to form cell-mats for analysis.

For the zeta potential measurements, rectangular shaped (W × L × H = 1 cm × 2 cm × 0.1 cm) solid materials were clamped to create a channel of 25 mm length and 5 mm width, with active layers facing each other and the charge measured (in mV) when growth medium (pH 7.0) as an electrolyte flowed through the channel. The zeta potential results were calculated using the Helmholtz–Smoluchowski equation.

Contact angles were measured using the sessile drop method on materials previously dried for 24 h at room temperature (25 °C). All contact angle values were obtained from at least six measurements using growth medium (100 μL) as solvent.

**Cytometry and microscopy**

Cell counts for the biofilms for each sample were measured following suspension of the biofilm material in sterile buffer solution, and normalized total number of cells to the surface area of the anode used for extraction. Bacterial cell counts were measured by flow cytometry (BD Accuri C6 flow cytometer, BD Biosciences, Franklin Lakes, NJ). Samples (700 μL) were transferred to a sterile Eppendorf tube and incubated at 35 °C for 10 min prior to staining with SYBR Green I (7 μL of 100× stock solution in 700 μL sample), vortexed, and then incubated again at 35 °C for 10 min. Samples (200 μL) were then transferred to a 96-well plate for cell counting.

Samples on electrodes were prepared for scanning electron microscopy (SEM) by placing the electrode in solutions of 2% glutaraldehyde containing 10 mM 2-[4-(2-hydroxyethyl)-1-piperazinyl] ethanesulfonic acid (HEPES, pH 7.4) buffer overnight at 4 °C and then 1% osmium tetroxide overnight, with washing using 10 mM HEPES buffer (pH 7.4) between steps (all Sigma-Aldrich). The samples were then dehydrated (5 min for each step) in a graded series of aqueous ethanol solutions (10−100%) and oven-dried (2 h at 40 °C) to remove residual moisture. The dried samples were mounted over SEM stubs with double-sided conductivity tape. After sputter-coating the samples with gold-palladium for 30 s at 25 mA current in an argon atmosphere (Emiteck, K550), SEM imaging (Quanta 200D, FEI, The Netherlands) was performed using an accelerating voltage of 25 kV and working distance of 10 mm.

For confocal laser scanning microscopy (CLSM) analysis, electrode sections were transferred into sterile vessels containing 50 mL of anaerobic acetate-free growth medium. The graphite rod was cross-sectioned into pieces (∼3 mm in height) using a scalpel and stained, by incubation in the dark for 15 min in 10 mL of 10 mM potassium phosphate buffer, pH 7.0, containing 1 μL of propidium iodide and 1 μL of Syto 9 from Molecular Probes BacLight LIVE/DEAD L7012 stain kit (Invitrogen Corp., Carlsbad, CA). The samples were then gently washed in phosphate buffer (10 mM, pH 7.0) to remove unbound residual dye from the biofilm matrix. The biofilm stained electrodes were placed on a multiwell microscope slide to examine thickness of biofilm through a Zeiss LSM 510 Axiovert inverted confocal microscope with a 40× Achroplan oil immersion lens. A minimum of 10 fields of biofilm views were imaged, and Z-series images were processed and analyzed with Zeiss LSM510 operating software for biofilm thickness measurements. Images were obtained using an excitation wavelength of 488 nm and a BP500−550 emission filter for green fluorescence. The excitation wavelength was 543 nm, and emission filter LP605 was used to obtain images for red fluorescence.

**Table S1: Surface characteristics of electrodes (n=2) using growth medium as solvent/electrolyte.**

| **Electrode** | **Zeta potential (mV)** | **Contact angle** |
| --- | --- | --- |
| -NH_2_ | -68 ± 6 | 89.2 ± 0.3 |
| -COOH | -56 ± 2 | 14.0 ± 0.5 |
| -OH | -37 ± 2 | 74.3 ± 0.7 |
| -C_2_H_5_ | -109 ± 2 | 99 ± 2 |
| Bare | -104 ± 3 | 104 ± 4 |
| *G. sulfurreducens* cells | -29 ± 5 | 30 ± 5 |
| *D. acetexigens* cells | -16 ± 4 | 20 ± 3 |

**Fig. S1. Voltammetric response of graphite rods during the functionalization of anode.**

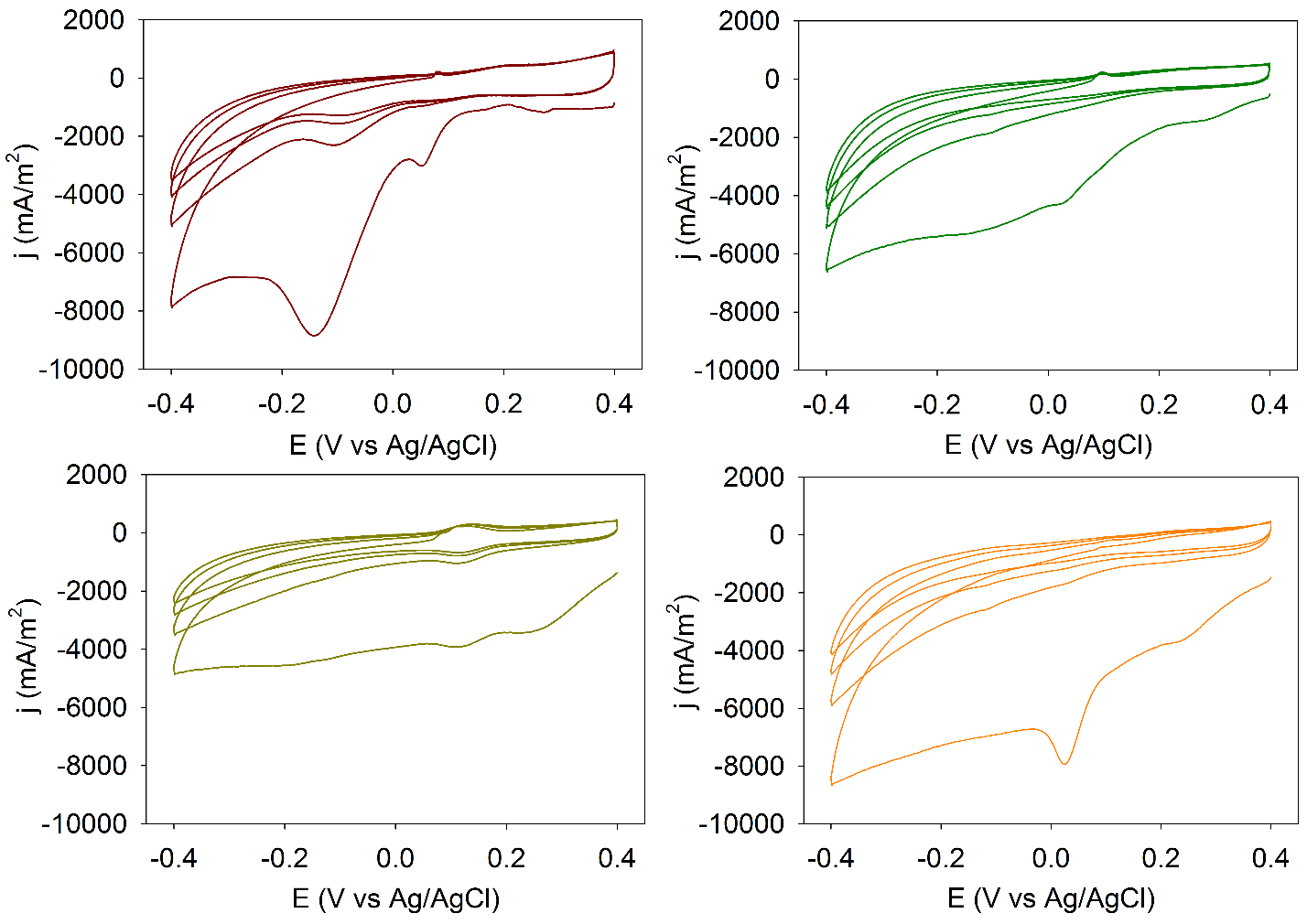

NH_2_

OH

COOH

C_2_H_5_

**Fig. S2:** **Amperometric response of matured biofilms under continuous flow feed at an applied potential of -0.1 V vs. Ag/AgCl**. Black arrows indicate the time when feed flow was switched from 0.5 L/day to 1 L/day, and asterisks represent the point when fed flow switched back to 0.5 L/day. Grey arrow indicates the time when the biofilms were sampled for SEM and microbial community analysis. Numbers represent the time where CV analysis was conducted.

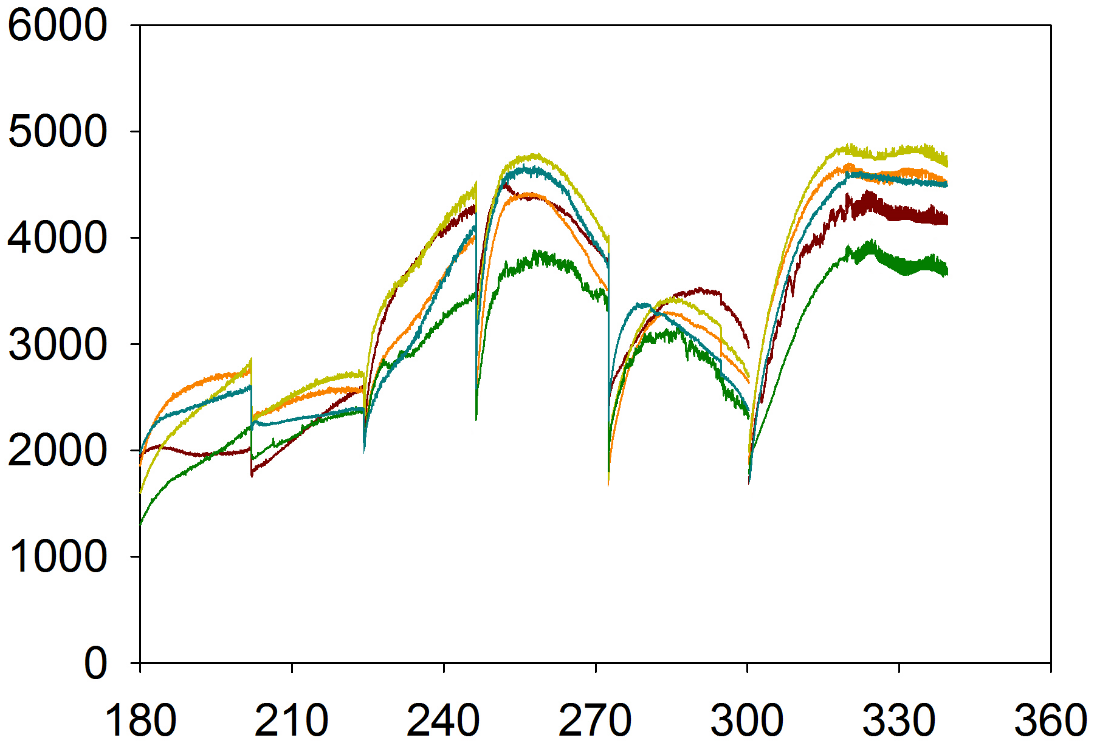

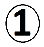

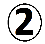

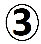

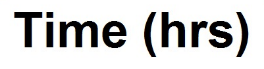

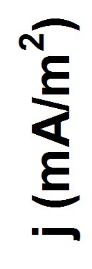

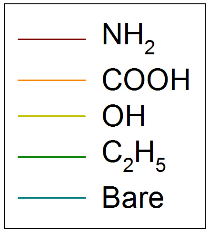

**Fig. S3. CV (1 mV/s) of biofilm extract.** Redox response confirms the presence of soluble mediators in the biofilm matrix.

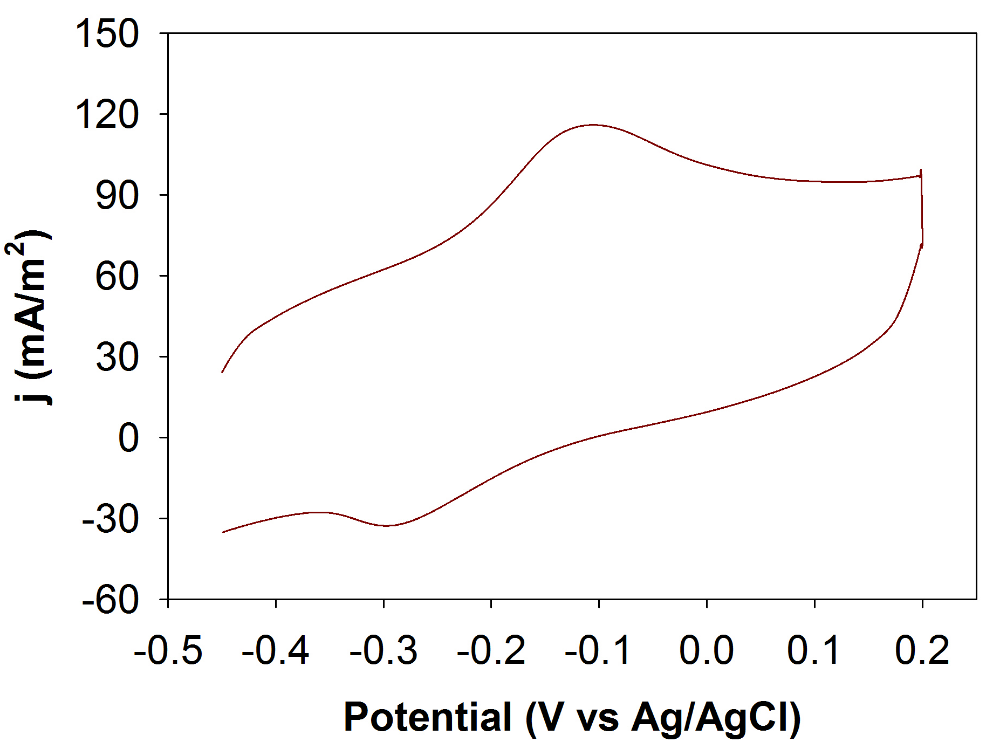

**Fig. S4. CLSM images of biofilms.** Thickness of matured mixed culture biofilms (340 h aged) grown on different anodes (unmodified and functionalized graphite electrodes) at an applied potential of -0.1 V vs. Ag/AgCl.

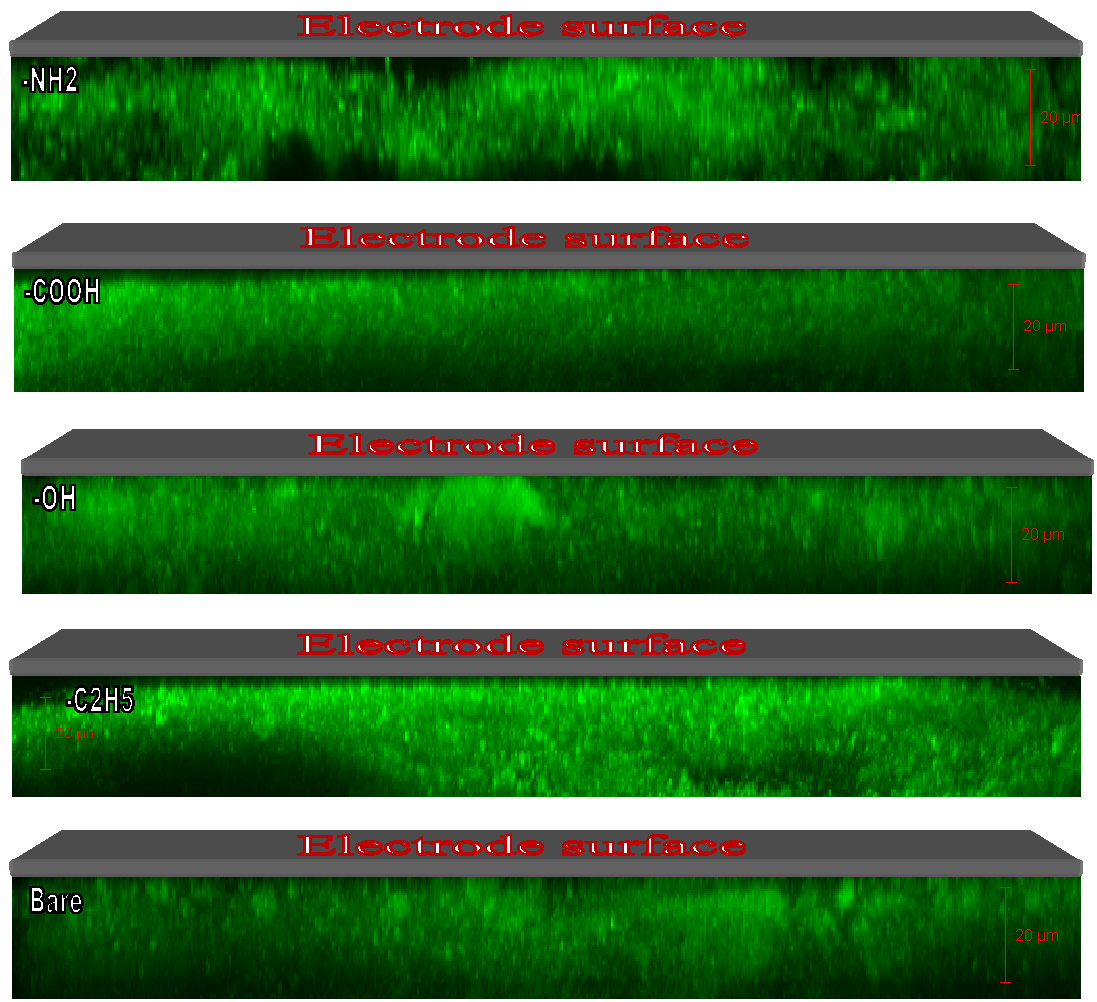

21 ± 2 µm

22 ± 2 µm

23 ± 4 µm

21 ± 4 µm

23 ± 3 µm

Fig. S5. Amperometric response of *D. acetexigens* on functionalized graphite electrodes at an applied potential of -0.1 V vs Ag/AgCl for initial start-up of *D. acetexigens* growth, in the presence of *D. acetexigens* growth medium containing 10 mM acetate as an electron donor and 10% w/v inoculum.

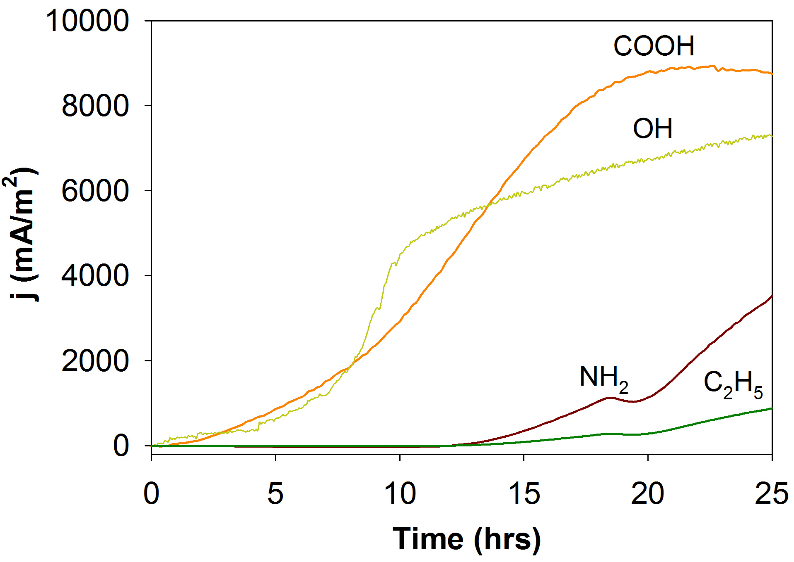

Preliminary tests were conducted to investigate the effect of surface chemistry on the growth of biofilms from pure cultures of *D. acetexigens* during the early phase of biofilm formation (Fig. S5) under -0.1 V vs Ag/AgCl applied potential control. Biomass and current density measurements indicate that electroactive biofilms of *D. acetexigens* are formed at all electrodes, but with a more rapid onset of current observed for electrodes functionalized to introduce -COOH, -OH and to a lesser extent -NH_2_ terminal groups compared to electrodes modified with hydrophobic -C_2_H_5_ groups (Fig. S5), similar to the trend observed for early-stage growth of biofilms from the mixed culture inoculum (Fig. 1A). A maximum current density of ~ 9 A/m^2^ is obtained less than 20 h after inoculation for the graphite electrodes with -COOH terminal groups, similar to that recorded for the first batch-feed at the unmodified electrodes (Fig. 5A). This suggests that the unmodified electrode, unlike the case with the mixed-culture inoculum, is a suitable surface for rapid start-up of early-stage *D. acetexigens* biofilm growth and performs like the -COOH surface.
